# Abandoning the Quadrupole for Mass Spectrometry Fragmentation Analysis: Towards 1,000 Hz Speeds with 100% Ion Utilization Using High Resolution Ion Mobility Precursor Isolation

**DOI:** 10.1101/2024.10.18.619158

**Authors:** Daniel DeBord, Leonard C. Rorrer, Liulin Deng, Frederick G. Strathmann

## Abstract

In today’s fast-evolving landscape of omics research, tandem mass spectrometry has become a cornerstone for uncovering the complexities of biological systems. Yet, despite its essential role, the technology remains bound by the inherent limitations of quadrupole filters, which throw away up to 99% of the useful ion signal and caps the speed at which fragmentation data can be generated. These deficiencies often force compromises in data depth and accuracy, hindering breakthroughs that tie genomics, proteomics, and metabolomics together. A new technology is poised to break through these performance barriers, unlocking unprecedented capabilities in precision, sensitivity and throughput. This step-change in how mass spectrometry fragmentation analysis is performed will reshape the future of scientific discovery, pushing the boundaries of what’s possible. The solution is high resolution ion mobility (HRIM), which offers a means to quickly and efficiently isolate ions prior to fragmentation and detection by a high resolution mass spectrometer (HRMS) while also resolving challenging isomeric and isobaric compounds that lead to chimeric MS/MS spectra. HRIM isolates ions in time as a result of a high speed separation rather than acting as a filter that discards ion signal like the pervasive quadrupole mass analyzer, allowing higher sensitivity analysis to be achieved. Also, since HRIM eliminates the need to hop or sweep electronics control parameters, as is the case with a quadrupole, fragmentation spectral generation can occur at a much faster rate, upwards of 500 Hz. This whitepaper describes an IM/MS methodology first contemplated over twenty years ago and today being positioned as the fastest method for high resolution fragmentation analysis. Revisiting this concept using the latest generation ion mobility technology based on structures for lossless ion manipulation (SLIM), which is the only HRIM technology that delivers similar resolution and range of analysis as a quadrupole, realizes the full potential of this approach to deliver benefits in both speed and sensitivity for high performance MS/MS measurements. This new way of achieving ion fragmentation in complex samples is set to revolutionize the mass spectrometry space, starting in the growing field of proteomics where all researchers are seeking faster methods to achieve more comprehensive proteomic coverage. Due to the advantages of HRIM physics, we predict this will set the bar for high throughput -omics within the coming years and will eventually be as ubiquitous as the quadrupole is today.

## The Problem with MS/MS

High performance mass spectrometry analysis is almost universally reliant on performing two or more stages of mass analysis with a fragmentation step between. This approach of tandem mass spectrometry, termed “MS/MS”, has proven to be the most reliable means to achieve a high degree of specificity about which molecules are being detected from the sample.^1^ It enables confident identifications and sensitive quantitation by removing signal associated with compounds that have the same precursor *m/z* but different fragment ions. However, common implementations of this methodology have a major drawback in that while a certain type of ion is being selected for fragmentation analysis, all other ions coming from the ion source are discarded, since they cannot be analyzed simultaneously. This sequential analysis requirement translates to an increasing loss in sensitivity and potentially missed compound identifications as the number of desired isolation targets grows, because the time spent analyzing each precursor ion (i.e. the dwell time or accumulation time) must be distributed across all targets of interest.

Researchers have traditionally focused on the duty cycle of an instrument as a key figure of merit to define its efficiency of operation, where duty cycle is defined as the percentage of time during an analysis cycle that an instrument spends recording MS1 (precursor) or MS2 (fragment) spectra. This metric emphasizes minimizing timing delays due to processes like isolation transitions or data transfer so that most of the experiment time is spent collecting data. Many modern instruments boast duty cycles that are 95-99%,^2-5^ but this metric can be misleading because it does not account for what fraction of the ions created in the source actually get detected. In other words, the instrument may be collecting data 95-99% of the time, but it may fail to detect certain key analytes because they are low abundance and require longer dwell times to be detected. For this purpose, we should think about an ion utilization efficiency (IUE) metric, which instead measures the percentage of ions of interest created by the ion source that are analyzed by the system and have a chance to make it to the detector. This IUE metric is a measure of how wasteful or efficient the mass spectrometry analysis methodology is with the ions being generated. When assessed in this way, many “high performance” MS systems are actually using less than 20% of the available ion signal, with some reaching to less than 1%!^6, 7^ This means upwards of 99% of potential ion signal is being discarded and will never reach the detector. While IUE may not be the most important variable for all MS experiments, for applications such as single cell proteomics or immunopeptidomics where strict sample limits exist, lower relative IUE creates strict limits on sample completeness, meaning that elusive, low abundance biomarkers remain just out of reach.^8^ In personalized medicine where the dream is to develop targeted approaches based on the immunopeptide profile of a tiny biopsy punch, increased relative IUE may be the difference between a dream and a medical reality.^9^

This trade-off between the sensitivity of detection for certain analytes and the total number of analytes that a system can analyze by MS/MS has propelled development of a wide array of data-dependent and data-independent acquisition (i.e. DDA and DIA) techniques.^10, 11^ It is well known that DDA is capable of generating higher quality MS2 spectra for a fewer number of targets and DIA can generate fragmentation spectra for “all” precursors in a reproducible way, albeit with the consequence of requiring software to untangle the chimeric MS2 spectra recorded. In the field of proteomics, this has led to a blurring of the lines between DDA and DIA in the form of narrow window DIA methods that purport to deliver the benefits of both: full precursor coverage and sufficiently narrow quadrupole isolation windows that limit MS2 spectral complexity.^12^ However, this approach makes an enormous concession in terms of the ion utilization efficiency, because narrower windows require more windows to cover a given *m/z* range. If a researcher configures their latest generation LC-MS system to isolate 200 windows (at a blistering speed of 200 Hz), the selected precursor(s) are only being analyzed for 5 ms out of every second it is present (0.5% ion utilization).^12^ This corresponds to a loss of 99.5% of the available ion signal and a 100x sacrifice in the measure signal to noise of fragmentation spectra.^12^ Similarly poor IUE is also observed for methods where the quadrupole isolation window is swept rather than hopped, also resulting in a reduction or limitation in the number of analytes that are detected.^13-16^

## Overview of How Different HRMS Instruments Generate MS/MS Spectra

Figure 1 demonstrates diagrammatically the steps in an MS/MS analysis cycle for various instrumentation approaches. In each case, a continuous beam of ions is introduced from the ion source and ion mobility, a quadrupole mass filtering, or both are used to isolate incoming precursor ions.

**Figure 1.**
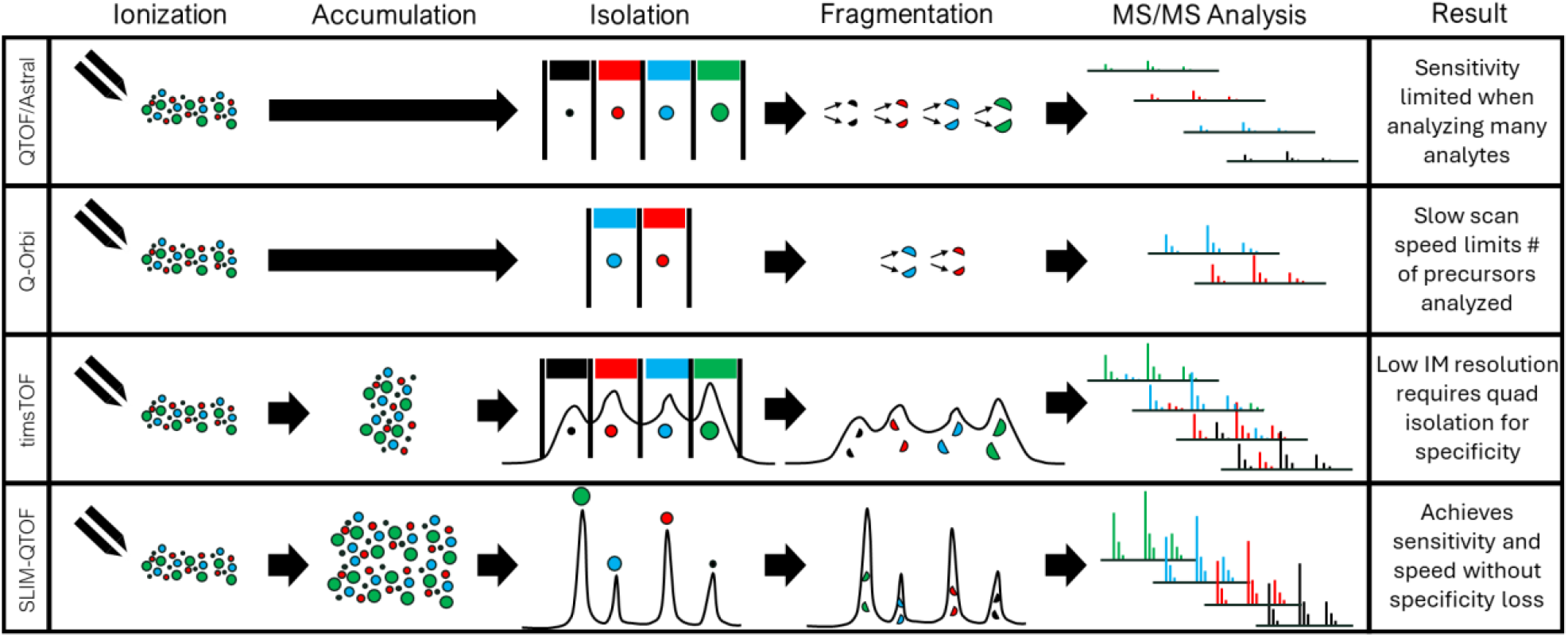
Diagram comparing various methodologies for generating fragmentation spectra.

## QTOF (and Astral) Instruments

For a typical QTOF (and Astral) mass spectrometer, the quadrupole mass filter removes all other ions from the beam other than the targeted precursor range during each dwell period. The signal intensity of the resulting MS/MS scan is therefore proportional to the time spent accumulating signal for that given precursor. Additional precursor isolations are performed sequentially, dividing the total time available per analysis cycle based on the total number of precursors to be analyzed. For samples with few coeluting precursors, this approach works well, but as sample complexity grows, sensitivity tends to fall off dramatically because less time is allocated to each precursor ion.

## Q-Orbitrap Instruments

In the case of a Q-Orbitrap instrument, field asymmetric ion mobility spectrometry (FAIMS) is often used to prefilter ions based on charge state, and for proteomics, to remove singly charged peptide ions. This allows the system to make the most use of the ion trapping capacity of the analyzer by avoiding filling the trap with nontarget ions. By incorporating an accumulation step post-quadrupole isolation (e.g. in the C-trap or linear ion trap), the Orbitrap system is able to generate higher sensitivity spectra for the precursors of interest. However, the slower speed of the Orbitrap (or linear ion trap) relative to a TOF mass analyzer limits the number of MS/MS spectra that can be generated in a given period of time.^17^

In order to overcome the inefficiencies of a quadrupole precursor isolation step, users would prefer to have a technology capable of pre-sorting the precursor ions so that they could be sequentially fragmented and analyzed by the mass spectrometer without filtering out other precursor ions of interest. The wasteful nature of quadrupole-based MS/MS was first recognized and a solution proposed nearly 25 years ago by David Clemmer’s lab in an article where they noted: “*Initial m/z selection is an inherently inefficient process since other m/z ions from the sample are discarded. […] We introduce a method […] based on an ion mobility/time-of-flight mass spectrometry technique that allows a mixture of ions to be separated prior to being dispersed into a mass spectrometer for m/z analysis*.”^18^ They called this approach “Parallel CID” because fragment ions generated appeared at parallel drift times with their corresponding precursors, enabling straightforward association and direct generation of fragmentation spectra for all precursor ions of interest without requiring a quadrupole filtering step.^19-21^ This means that the isolation typically performed using a quadrupole could potentially be replaced with ion mobility. Per Clemmer’s pioneering work, “*This [ion mobility] time scale is ideal for feeding complex mixtures into high-speed mass spectrometers [such as time-of-flight (TOF) instruments] in a high-throughput fashion*.”^21^ However, the authors noted that the applicability of this approach for complex mixtures was limited by the resolution of their mobility analyzer (20-30). Other researchers and instrument developers have attempted this mode of operation with varying degrees of success,^22, 23^ but it would take over 15 years for a new generation of higher performance mobility analyzers to be introduced that would reveal the true potential of Parallel CID.

## timsTOF Instruments

As shown in Figure 1, the Bruker timsTOF system^2^ leverages Clemmer’s parallel CID methodology, but also introduces the ability to accumulate ions while performing mobility separation of a previous packet of ions. This allows IUE to be further improved, even beyond what Clemmer had posited. However, the peak capacity of the trapped ion mobility spectrometry (TIMS) analyzer is still restricted to approximately 20 when analyzing the full mobility range, corresponding to 5 ms wide peaks within a typical 100 ms separation timescale for each scan, which caps the maximum speed and specificity of precursor isolation.^15^ The ion mobility separation results in precursor ion packets that are concentrated in time and queued up for isolation by the quadrupole. But the limited peak capacity means that quadrupole isolation is still required for most complex samples in order to achieve sufficiently clean MS2 spectra to allow for identification. The quadrupole can be hopped or swept across the mass range to align isolation timing with the desired precursor ion peaks so that fewer target ions are discarded in the isolation process. This approach has become known as parallel accumulation with serial fragmentation (PASEF), with many iterations of the implementation being developed to continuously improve upon the duty cycle of the device. However, because of the low resolving power and low peak capacity in the mobility dimension, there remains a need for narrow quadrupole isolation to create useful MS2 spectra, meaning the TIMS separation scan must be performed multiple times so that the isolation window can be stepped or ramped across the full range of possible *m/z*’s at a given mobility. The need to cycle through multiple TIMS ramps results in an ion utilization tradeoff such that the number of TIMS scans with quadrupole isolation determines the period of time over which certain precursor ions can be successfully transmitted for detection, while during all other TIMS scans of a given cycle those precursor ions are discarded due to the filtering effect of the quadrupole.

## A Faster and More Sensitive Way to Generate Fragmentation Spectra

As indicated by Clemmer’s early work, in order to overcome the need for a filtering step when analyzing complex mixtures, a higher resolution ion mobility approach is required. For reference, typical DIA workflows employ quadrupole isolation windows of 2-20 Th wide. At 600 Th, this corresponds to a resolving power of 30-300 (M/ΔM), setting the bar for the level of specificity required. In 2015, a suitable high resolution ion mobility technology was introduced in the form of Structures for Lossless Ion Manipulation (SLIM).^24-26^ The SLIM technology enables significantly longer ion mobility path lengths (e.g. 13 m) in a reasonably small size, owing to its printed circuit board-based electrode design that enables the ion path to be folded back on itself in a serpentine arrangement. This results in ion mobility resolving powers of 250-300 (CCS/ΔCCS),^27^ making it the only ion mobility technology that can provide sufficiently high resolving power over the full mobility range in a single scan to enable direct precursor isolation without requiring quadrupole filtering for complex samples.

The other significant advantage of the SLIM technology is that it allows for much larger ion populations to be accumulated and stored, with capacities of over 10^8^ charges having been demonstrated, which is 10-100x higher than TIMS and Orbitrap devices.^28^ Greater ion capacity results in the ability to increase the sensitivity and dynamic range of analysis and ensures that maximum ion utilization through parallel accumulation can be realized, even for parallel separation time frames of 100’s of milliseconds.

## The SLIM-QTOF Approach to Optimize Ion Utilization and Achieve Quad-Free Fragmentation Analysis

A technique termed mobility aligned fragmentation (MAF), which is somewhat analogous to Clemmer’s Parallel CID, is displayed diagrammatically in Figure 2. In this mode, alternating ion mobility-mass spectrometry data frames are recorded without (MS1) and with (MS2) fragmentation energy applied via a collision cell prior to recording a full mass spectrum for every mobility time point using a TOF analyzer. This results in MS1 data frames which capture the *m/z* and arrival time features for all precursor ions present without having to filter out ions with a quadrupole, thereby compromising speed and sensitivity of analysis. The MS2 frames display unique *m/z* peaks for all fragment ions generated from all precursor ions, isolated via the mobility peak profiles which are consistent with their respective precursors in the MS1 frame.

**Figure 2.**
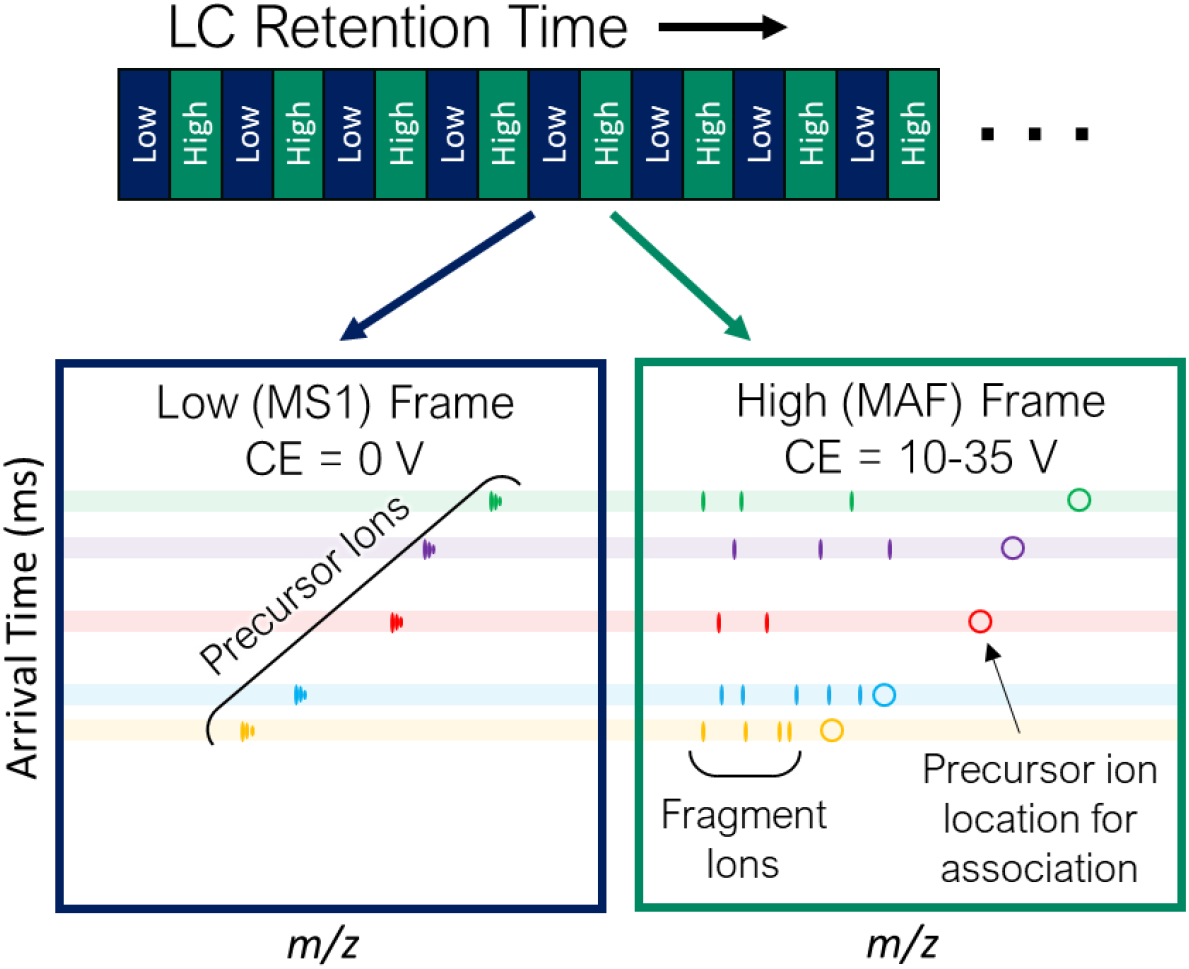
Diagram showing the approach for generating Mobility Aligned Fragmentation data, consisting of alternating Low and High collision energy (CE) data frames which contain precursor and fragment ion spectra, respectively. Association of precursors and fragments is achieved based on arrival time alignment in the ion mobility dimension.

In order to demonstrate the potential of a SLIM-QTOF instrument to achieve this type of high-speed, “quad-less” HRIM/MS operation, a commercial MOBIE HRIM system (MOBILion Systems, Inc.) with a 13 m ion mobility path length coupled to an Agilent 6546 QTOF was used to generate a proof-of-concept MAF data set by infusing a dilute mixture of 15 peptides^1^ and ionizing by nano-electrospray. Figure 3 demonstrates the correlation between extracted ion mobiligrams for all 15 peptides with precursor ions (MS1) in panel A and the most abundant fragment ions (MS2) for each precursor displayed inverted in panel B. As indicated by the similarities in HRIM arrival time and peak shape between each precursor and fragment ion, association of the fragment ions is intuitive and entirely analogous to the precursor/fragment association algorithms currently employed in chromatography.^29, 30^

**Figure 3.**
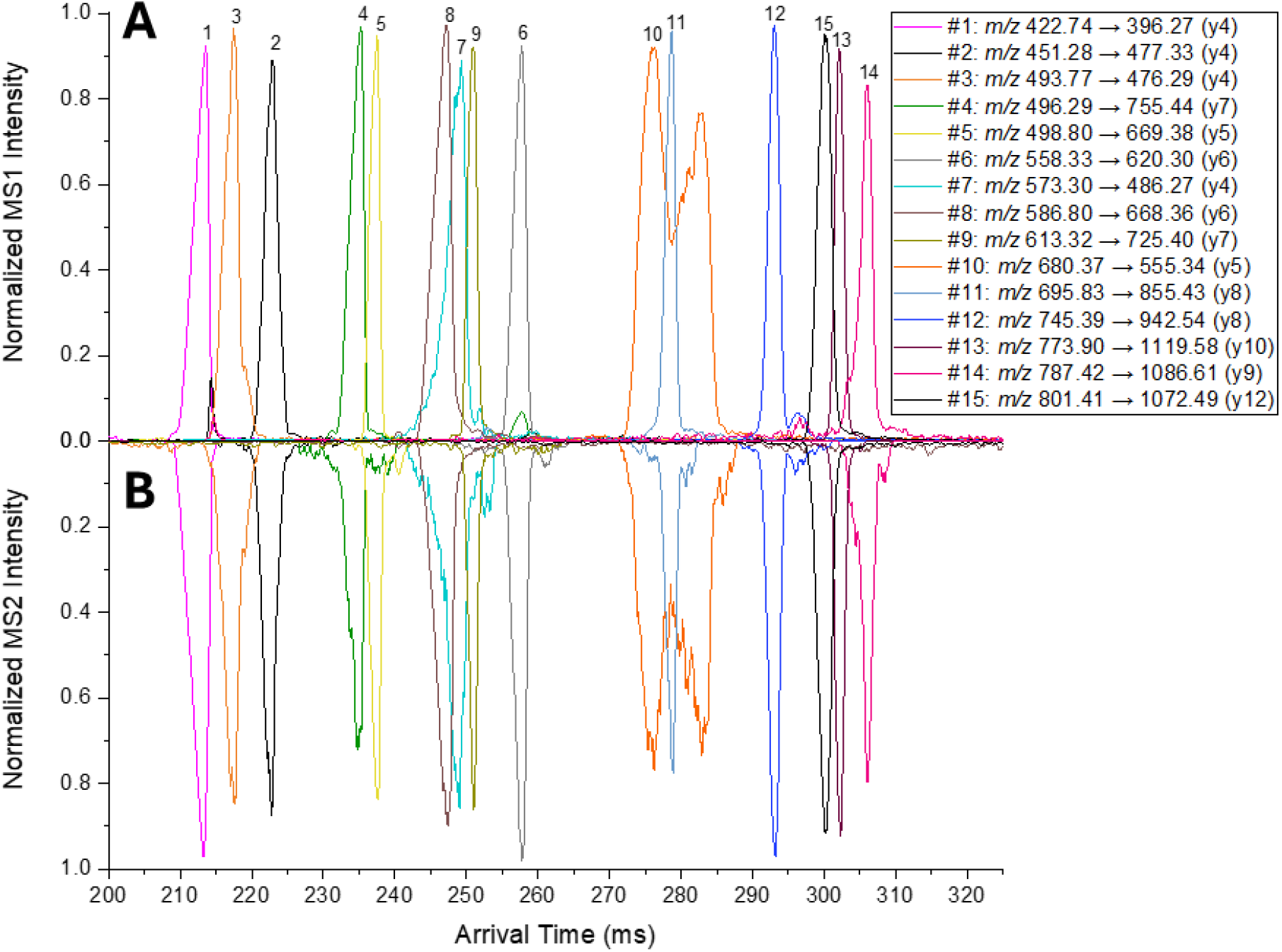
Extracted ion mobiligrams for 15 peptides listed in Table 1. **(A)** precursor ions from MS1 data frames and **(B)** base peak fragment ions from MS2 data frames demonstrating mobility alignment of the precursors and fragment ions.

The observed mobility peak shapes and widths are somewhat variable, depending on the signal abundance and conformational stability of each peptide, with well-behaved ions typically exhibiting peak widths (at half maximum) on the order of 1.5 to 2.5 ms. The asymmetry and variability of the peak distributions for certain peptides provide an additional identifier that enhances one’s ability to deconvolve overlapping features and better determine which fragments came from which precursor mobility peak. In this simple example, we demonstrate the ability to generate fragmentation spectra for all 15 peptides in a period of just 100 ms, directly demonstrating 150 Hz MS/MS. However, as indicated by the empty regions of the spectrum, the peak capacity in the separation domain is sufficient to triple or quadruple the number of components analyzed while still allowing for accurate extraction of unique fragmentation spectra. Also, this particular mixture only contains doubly charged peptide ions up to ∼1600 amu in size. Allowing for larger peptides and other charge states, which are typically observed for more complex proteomic digests, expands the mobility range of analysis by 1.5-2x in the arrival time dimension. Based on this analysis, if we assume a typical mobility peak width of 2 ms per peptide and a separation time scale of 200 ms, this type of 13 m HRIM MAF experiment is capable of generating fragmentation spectra at upwards of 500 Hz. Allowing for deconvolution to extract partially overlapping features (as is already done for chromatographically coeluting compounds), it is easy to imagine that HRIM/MS spectra can be generated at a rate of more than 1,000 per second. Most importantly this speed can be achieved without sacrificing sensitivity because ions are accumulated in parallel without loss or filtering. By accumulating ions for a period equal to the separation period, ∼100% ion utilization efficiency can be realized. The secondary benefit of this approach is that low abundance analytes experience a pre-concentration effect, with 100’s of ms worth of ions being focused into a narrow ion mobility peak that raises the signal level above the typical noise floor of the instrument, thereby enabling detection. This does challenge the detection system to handle the high dynamic range of signal observed per TOF transient for abundant species, but modern high speed digitizers and electron multiplier detectors are available to address this need, with current systems being capable of achieving >4 orders dynamic range for a single mobility frame of data.

**Table 1.**
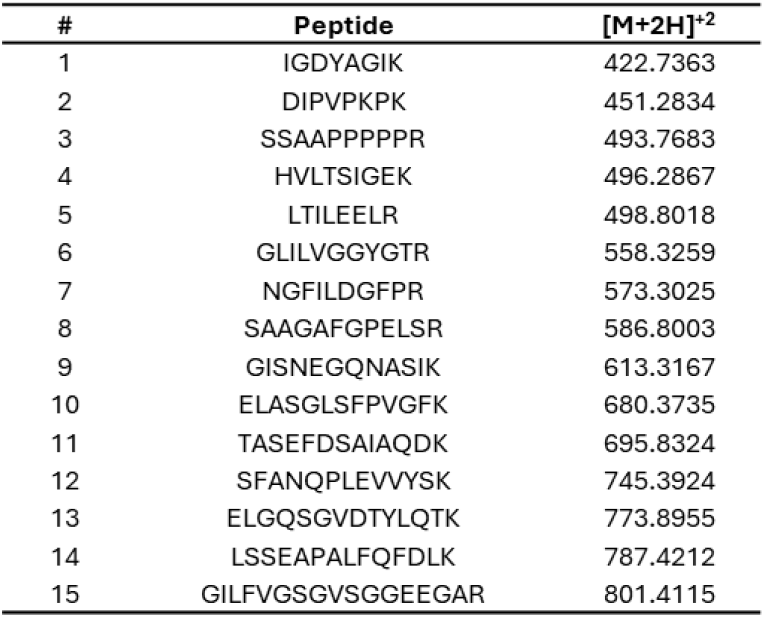
List of peptide sequences and corresponding accurate masses for the doubly charged ions observed from the Pierce peptide retention time calibration mixture.

## Parallel Accumulation with Mobility Aligned Fragmentation Outperforms Traditional MS/MS

To demonstrate how parallel accumulation with mobility aligned fragmentation (PAMAF) boosts the signal level for low abundance analytes, we compared MS1 and MS2 data collected on the same system as above, operating the SLIM component so that no accumulation or mobility separation was performed and ions were simply passed through the SLIM ion guide for QTOF analysis. Figure 4A shows the precursor mass spectrum for the peptide mixture corresponding to 5 ms worth of dwell time (equivalent to a 200 Hz acquisition rate). The signal intensity and spectral quality is quite low given that a very limited number of ions are transmitted during this short time period for such a dilute sample. Figure 4C shows that isolating a given precursor ion (*m/z* 801.2 corresponding to the doubly protonated precursor of GILFVGSGVSGGEEGAR) and fragmenting within the CID cell of the QTOF results in a very low quality fragment spectrum, with certain expected fragment ions for the peptide barely detectable. However, using a MAF approach with a 500 ms ion accumulation time prior to HRIM-MS detection, a ∼100x increase in signal levels for all precursor ions in the MS1 spectrum can be achieved, as shown in Figure 4B. This increase is due to SLIM’s ability to store up the ion signal for all incoming ions over 500 ms and then deliver separated mobility peaks just 2 ms wide to the QTOF for detection. As expected, Figure 4D shows that by simply applying a collision energy to all ions that elute during the mobility separation period, a ∼100x increase in spectral intensity for fragment ions can also be realized. Since no duty cycle limitation of the quadrupole is incurred, similar boosts in fragmentation spectra for all 15 peptides can be achieved.

**Figure 4.**
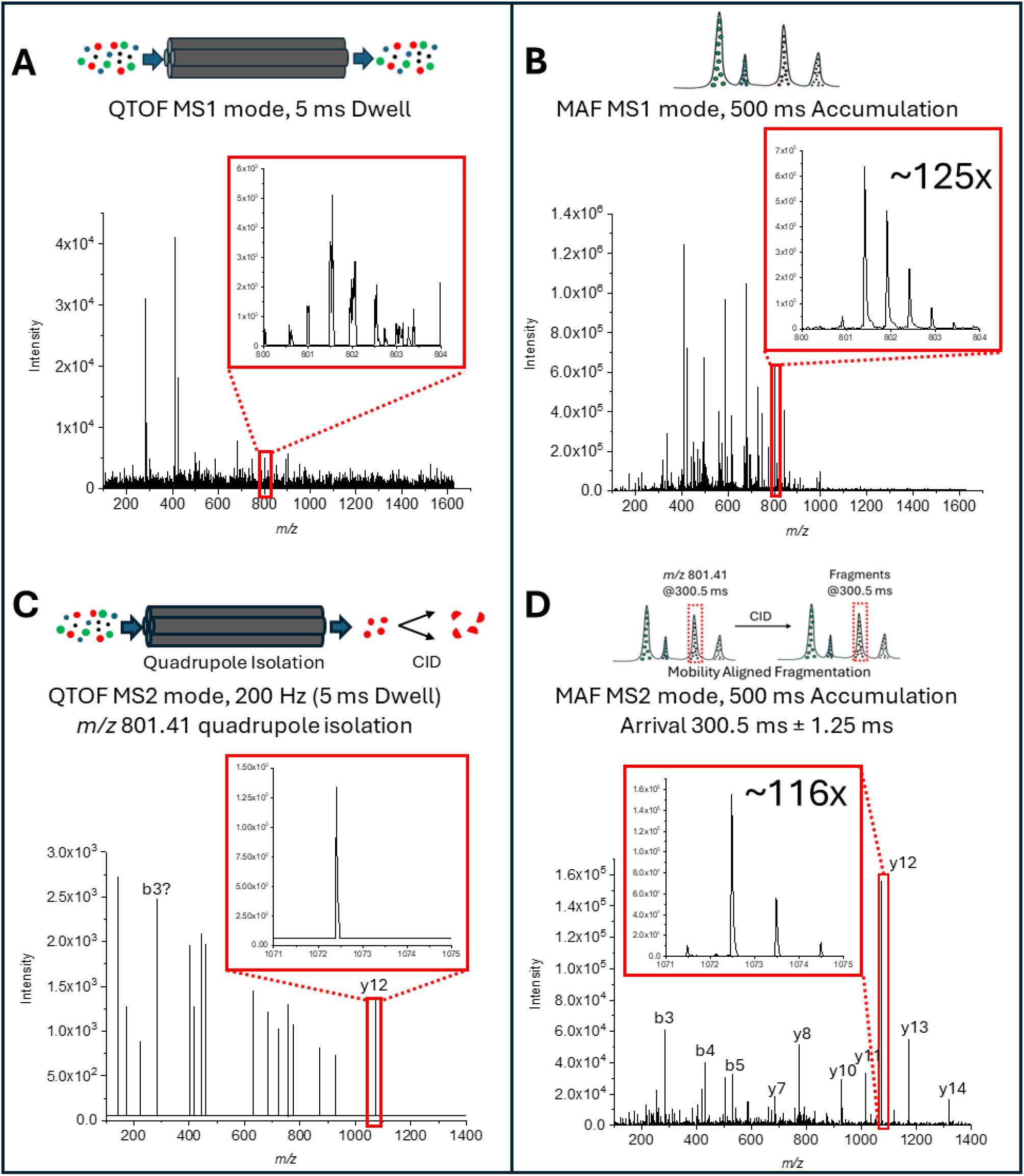
Comparison of MS1 spectra generated for the dilute Pierce peptide retention time calibration mixture in **(A)** QTOF mode and **(B)** MAF mode. Fragmentation spectra for the peptide GILFVGSGVSGGEEGAR at a collision energy of 21 V collected in **(C)** QTOF mode equivalent to 200 Hz MS/MS and **(D)** MAF mode equivalent to >500 Hz MS/MS. MAF spectrum corresponds to 500 ms worth of ion signal but extracted from an arrival time range of 2 ms centered on the detected mobility peak. In each case, the MAF mode of operation achieves ∼100x higher signal level to the QTOF mode of operation.

**Figure 5.**
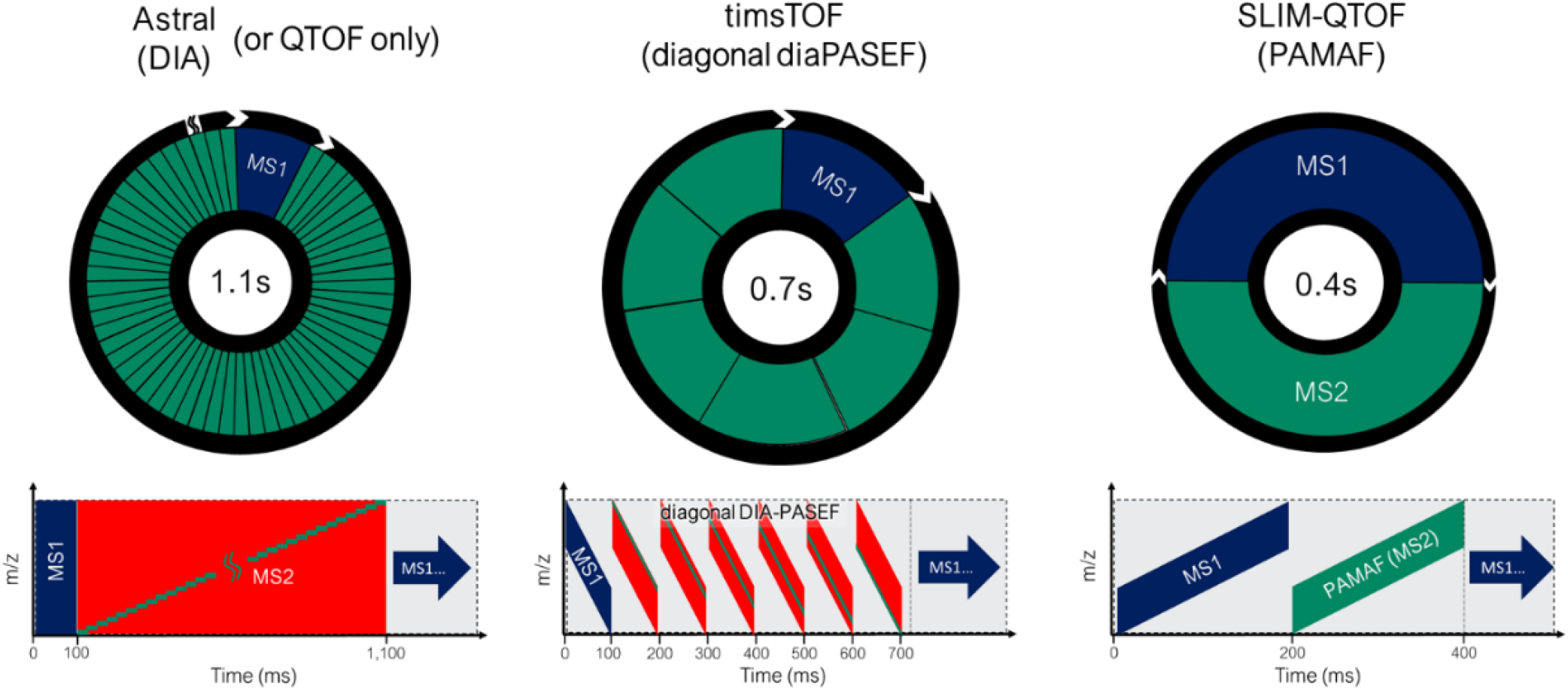
Comparison of analysis cycles for the Thermo Astral, Bruker timsTOF, and MOBILion Systems SLIM-QTOF systems.

## PAMAF Outperforms Current Best in Class MS/MS Systems

This new approach of rapidly fragmenting mobility peaks in real time without quadrupole isolation (PAMAF) has the potential to overtake current industry-leading LC-HRMS systems in terms of the speed and sensitivity of MS/MS measurement. To demonstrate this potential, we have generalized key performance metrics for the Thermo Astral and Bruker timsTOF instruments from relevant publications.^4, 12, 14, 15, 31^ By analyzing a typical analysis cycle of each instrument we can benchmark the proposed SLIM-QTOF methodology against these high-end instruments for a representative application in DIA proteomics. In the case of the Astral system, a 100 ms ion accumulation period is used to generate a full spectrum MS1 scan using the Orbitrap analyzer. While this MS1 mass measurement is occurring, 200 sequential ion packets are independently mass filtered with different quadrupole isolation ranges and then fragmented and analyzed via the Astral (TOF) analyzer. The high-speed and high transmission efficiency ion processing device allows up to 200 of these fragment spectra to be generated in just 1 second, however, this also means that each spectrum corresponds to <5 ms worth of ion signal. Per the ion utilization efficiency calculation introduced previously, this means that only 0.5% of each precursor ion introduced to the instrument are used to generate MS2 signal.^12^ The Astral system is able to overcome this high degree of inefficiency due to the highly optimized and efficient ion processing unit which collects and pulses ions into the TOF portion of the instrument.^4^ Still, this analysis demonstrates that an instrument that makes better use of the available ion signal has the potential to significantly improve the sensitivity of analysis. Also, the strict reliance on quadrupole isolation means that the MS/MS speed is capped at 200 Hz and that a full cycle takes over 1 second to complete, degrading quantitative accuracy for short LC gradients due to the limited number of data points that can be recorded across narrow LC peaks.

As noted above, the Bruker timsTOF system^2^ employing a dual TIMS cell configuration for parallel accumulation and ion mobility separation makes more efficient use of the ion signal available. A typical diagonal DIA-PASEF analysis cycle which balances sensitivity and speed can use a sequence of seven TIMS ion mobility ramps per analysis cycle. This includes a combination of one MS1 TIMS ramp followed by six MS2 TIMS ramps, where the quadrupole is used to sweep over a different range of precursor ions for each MS2 TIMS ramp. Each TIMS ramp corresponds to 100 ms for a total cycle time of 700 ms. The ion signal is divided among the seven different TIMS ramps, meaning that approximately 14% (1/7^th^) of a given precursor ion’s signal will be translated into a recorded fragment spectrum. It is critical to note however, that transmission efficiency through the TOF injection optics limit the overall sensitivity of the instrument, with estimates of the efficiency in the range of 5-25%.^6^ Since each mobility feature has a peak width at half height of approximately 5 ms,^15^ the maximum peak capacity per ramp is ∼20 and the maximum rate of fragment spectrum generation is 200 Hz. ^32^ This MS2 speed limit is also applicable to the timsTOF Ultra system, despite that system being specified at 300 Hz. In other words, the broad ion mobility peaks of the TIMS analyzer are too wide to make full use of the speed of the quadrupole electronics.

As shown in Table 2, we envision a SLIM-QTOF methodology based on mobility aligned fragmentation that would overcome the IUE and speed limitations outlined for these other systems. With SLIM, 2 ms wide ion packets can be generated for the full range of precursors over a time period of just 200 ms. An analysis cycle that includes a single MS1 scan followed by a single MS2 scan results in a cycle time of just 0.4 seconds, which is nearly 2x faster than the timsTOF and 3x faster than the Astral. By accumulating ions in parallel with each HRIM separation, 50% ions are used for MS1 and 50% of ions are used for MS2, meaning that ∼100% operational efficiency can be realized. Proprietary ion processing techniques to overcome the traditional limitations in ion transfer efficiency for TOF injection optics will mean that this boost in IUE is not sacrificed to losses in that region. Additionally, because precursor ion packets can be temporally focused by a factor of 100x and isolated by mobility arrival time, over 100 fragmentation spectra can be generated every 200 ms, resulting in an effective fragment spectrum generation rate of over 500 Hz. With algorithmic deconvolution of overlapping mobility features, we predict that >1,000 Hz HRIM/MS may be possible. In other words, deconvolution permits one to exceed the typical peak capacity limit to allow for more components to be analyzed within the same separation time period.

**Table 2.**
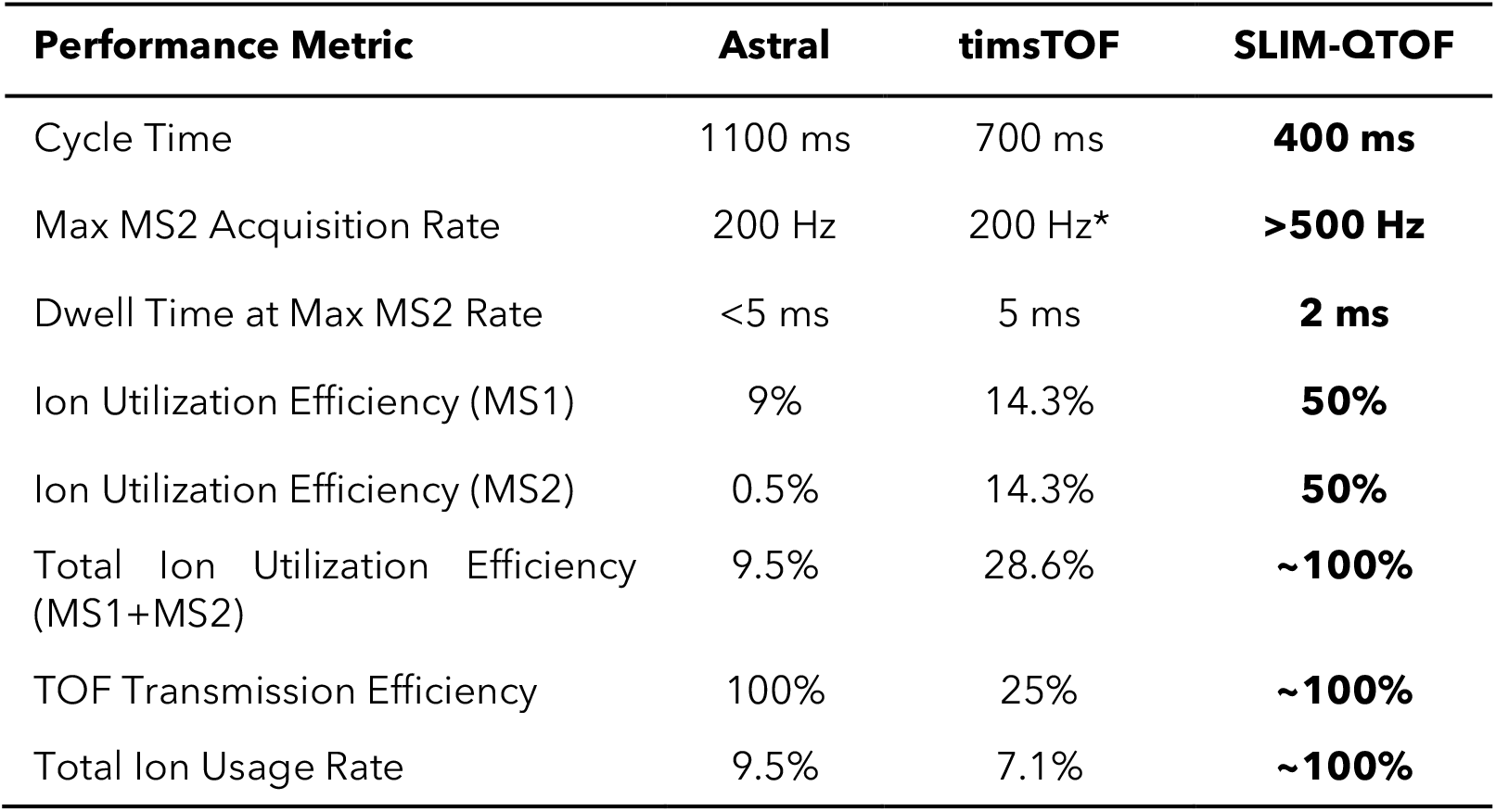
Comparison of various performance metrics between the Thermo Astral, Bruker timsTOF, and projected MOBILion SLIM-QTOF instruments. The estimates in the table are intended to be representative of typical performance but will vary for different acquisition method settings. *As noted in the text, the specified maximum rate of 300 Hz for certain timsTOF models is not achievable in practice.

Table 2 also calculates a total ion usage rate which is the product of the total ion utilization efficiency and the ion transmission efficiency. As noted above, the Orbitrap Astral system provides high speed and transmission efficiency but sacrifices sensitivity due to the duty cycle tradeoff of the quadrupole filter. The timsTOF system makes improvements in terms of ion utilization efficiency relative to the Astral, but sacrifices much of the signal to the lossy nature of TOF transfer optics.^6^ Interestingly, this comparison shows that similar levels of ultimate sensitivity and speed are achievable with both platforms which supports the numerous observations through bottom-up proteomics benchmarking studies that both latest generation platforms are roughly equivalent in terms of raw performance. By addressing the operational and transmission deficiencies of these platforms, the SLIM-QTOF approach offers an opportunity to realize an order of magnitude increase in sensitivity compared to the Astral and timsTOF.

## HRIM Uniquely Alleviates Chimeric Spectra and False Identifications

The value of HRIM/MS analysis extends beyond just the speed and sensitivity improvements described in the previous sections. The fact that MS/MS relies on low resolution mass isolation renders it completely blind to most isomeric and isobaric interferences that are universally present in complex samples. These compounds represent “known unknowns” that are largely ignored and instead manifest as low-quality spectral matches that prevent confident identification of convolved spectra. MAF is perfectly suited to tackle this challenge of isobaric and isomeric overlap of precursors that leads to chimeric fragmentation spectra. As shown in Figure 6,coeluting isobaric peptides can be physically resolved by HRIM prior to fragmentation, enabling extraction of purified fragmentation spectra. This is just one example of the thousands of coeluting isobars that were observed from this representative HeLa digest data set.^3^ As the desire to push omics analyses to ever shorter gradients takes hold, the need for a technique to address chromatographic overlap will become even more essential. The substantially higher mobility resolution of the SLIM approach (while operating over a full analysis range at speed) builds upon early attempts of the timsTOF platform. The ion mobility information encoded into the data can also be extracted via calibration of the mobility separation dimension. The progress toward leveraging collision cross section measurements (which quantify the size and shape of an ion) become even more powerful as the precision of the mobility measurement improves beyond 0.1%, as observed with HRIM.^33^ This means that an additional metric can be leveraged to confirm putative annotations and eliminate false identifications from being reported.

**Figure 6.**
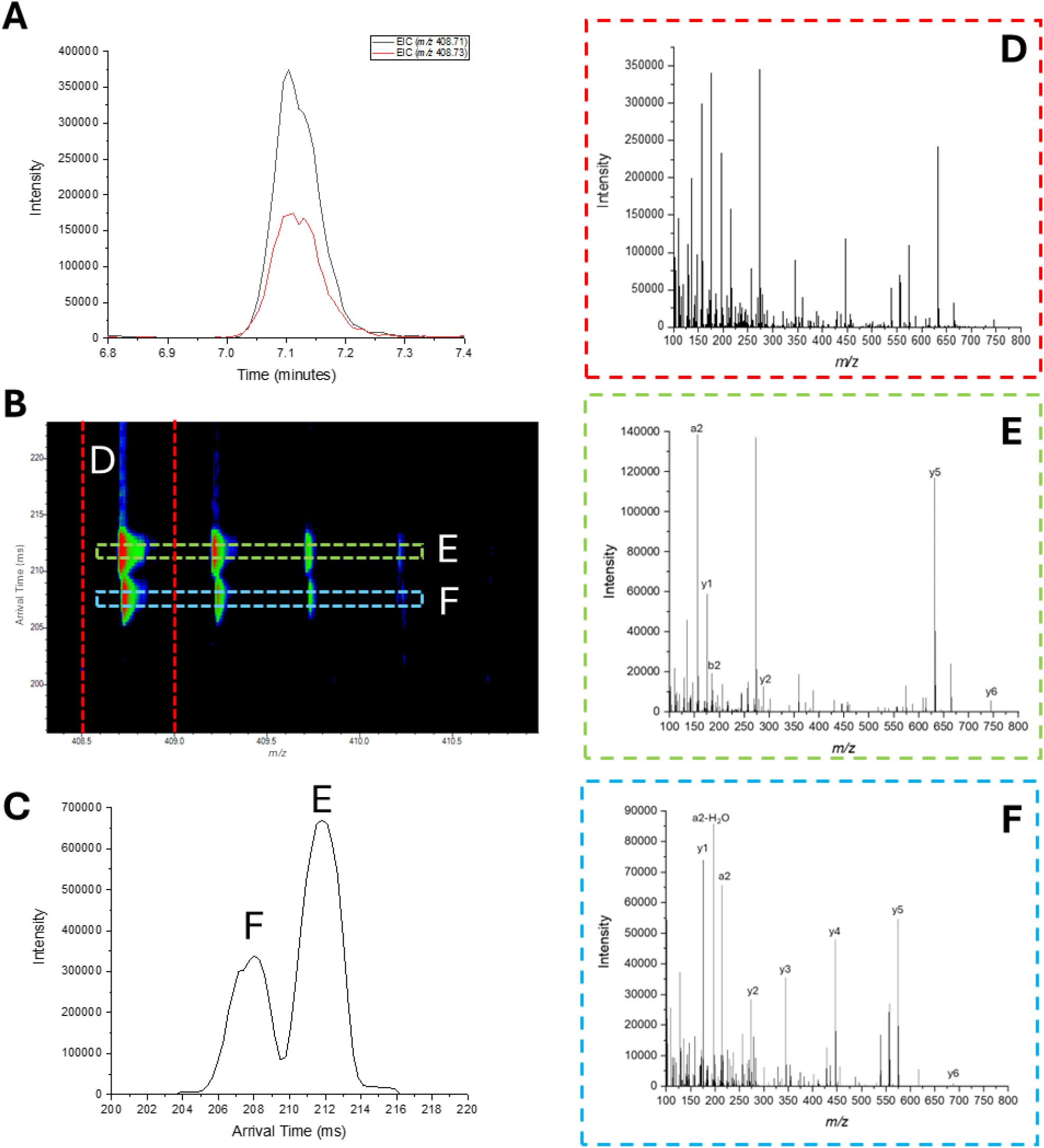
PAMAF prevents chimeric spectra from coeluting isobars. **(A)** Extracted ion chromatograms for coeluting isobaric peptides AIDEVNR (*m/z* 408.7141) and EIQTAVR (*m/z* 408.7323) from a representative proteomic mixture (100 ng HeLa) analyzed via MAF. **(B)** Mobility arrival time vs *m/z* heatmap plot and **(C)** extracted ion mobiligram for the two peptides demonstrating mobility-based separation. Extracted fragmentation mass spectra corresponding to isolation based on **(D)** the monoisotopic precursor *m/z* or **(E/F)** *m/z* and mobility precursor selections. Relying on *m/z* isolation alone as in D results in chimeric spectra while mobility filtering enables generation of cleaner fragment spectra.

## Conclusions

Analytical performance for the analysis of complex mixtures is ultimately a measure of how well an instrument can provide specificity without sacrificing sensitivity. The analysis provided in this whitepaper highlights key limitations in the traditional quad-based approach of using inherently low resolution filtering devices to achieve specificity. By instead leveraging a high-speed separations technology in the form of high resolution ion mobility, a more efficient and performant analyzer can be realized. Based on initial proof of concept data demonstrating the fundamental concepts of how PAMAF can be used to maximize ion utilization efficiency and speed of ion, we believe that it is possible to deliver a system that can generate >1,000 fragmentation spectra per second while achieving near lossless transmission and isolation of all precursor species. The steady progression of instrumentation development over the years has brought us to the point where we are within reach of maximum efficiency of operation. Building on past achievements limited by the technology of that time, we envision a future with ion mobility-enhanced fragmentation analysis that has been 25 years in the making. We anticipate that the resulting capabilities, unique to SLIM-based ion mobility, will enable untold scientific discoveries.

## Footnotes

1 Pierce Peptide Retention Time Calibration Standard, Thermo Fisher Scientific PN 88321, diluted to 100 fmol/μL

2 This description is generalized to represent performance achievable on all latest generation timsTOF models (Pro 2, HT, Ultra)

3 Pierce HeLa Protein Digest Standard, Thermo Fisher Scientific PN 88328, 100 ng injected on column using the Evosep 30SPD configuration

